# Inducible estrogen receptor alpha in normal breast epithelial cells demonstrate estrogen receptor-dependent DNA damage

**DOI:** 10.1101/2025.03.13.643100

**Authors:** Prabin Dhangada Majhi, Amye L. Black, Aman Sharma, Janhavi Phadkar, Amy L. Roberts, Puneet Singh, Jeffrey J. Kane, Kierney O’Dare, Patrick Van Eijk, Simon H. Reed, Sallie S. Schneider, D. Joseph Jerry

## Abstract

**Background:** Signaling by estrogen-receptor alpha (ERα) plays a major role in breast cancer initiation Investigations of the mechanism of DNA damage mediated by ERα signaling are carried out in breast cancer cell lines due to the lack of ERα+ normal human breast epithelial cells lines (HBEC). Defining the mechanisms by which ERα induces DNA damage and initiates tumorigenesis requires normal HBECs that express ERα, demonstrate estrogenic responses, and are amenable to long term propagation in culture.

**Methods:** We utilized lentiviral expression of an inducible ERα construct to generate four HBEC lines (HBEC-*ESR1*). We studied these cells for ERα-dependent responses using a luciferase reporter, endogenous gene expression and proliferation assays. RNA-Seq was performed to characterize the ERα-mediated transcriptomic patterns in the four HBEC lines. ERα mediated DNA double strand breaks (DSBs) were analyzed using γH2AX immunofluorescence.

**Results:** Expression and functional activation of ERα were observed in all HBEC-*ESR1* lines, whereas proliferation in response to 17β-estradiol (E2) was observed in 3 of the cell lines. Proliferative responses were due to intrinsic signaling within the HBECs as conditioned media from the cells failed to cause proliferation. A total of 682 genes were differentially expressed at 24h following treatment with 10nM E2 with 43% of these genes were also observed in ERα+ breast cancer cell lines (MCF7 or T47D). Gene-set enrichment analysis identified differential expression of genes in ERα signaling pathways and DNA repair pathways in E2-treated cells. E2-induced ERα signaling also increased γH2AX foci in 3 of the 4 cell lines. Levels of DSBs were increased by inhibition of the non-homologous end-joining (NHEJ) and homologous recombination (HR) pathways. DSBs were also increased in MCF10A-*ESR1* cells heterozygous for the *BRCA1*^185delAG^ mutation causing a truncated protein.

**Conclusions:** Inducible expression of ERα in immortalized HBECs recapitulate transcriptional, replicative and DNA damage responses. Increased DSBs in MCF10A-*ESR1* cells with heterozygous mutation of *BRCA1* indicate haploinsufficiency and the potential for increased genetic instability due to ERα signaling.

## INTRODUCTION

Breast development during puberty is driven by estrogens and is a critical window affecting lifelong breast cancer risk [1,2]. ER signaling can also cause DNA damage that contributes to acquired resistance to endocrine therapies. Analysis of mammary tissue shows that E2 causes DNA damage in mice but varied among strains [3,4]. E2-stimulated DNA damage has been attributed to 1) direct interaction of E2 to DNA [5,6], 2) ERα co-activator transcriptional complexes [7,8] and 3) replication stress induced by mitogenic E2 signaling [9,10]. Defining the mechanisms of ERα-mediated DNA damage are critical for establishing differences in breast cancer risk among individuals as well as risk of recurrence of breast cancers. The mechanisms of ERα mediated DNA damage have been studied primarily in ERα+ breast cancer cell lines such as T47D and MCF7 [7–10]. These breast cancer cells have undergone extensive genomic alternations including chromosomal instability and express higher levels of ERα than normal breast epithelium which has heterogenous levels of ERα expression [11–14]. These cell lines also lack the more than 300 genetic polymorphisms linked to breast cancer risk that may modify ER signaling [15]. For example, multiple studies have shown association of *FGFR2* risk alleles to ER+ breast cancer [16–20]. Guo et. al. conducted functional validation of 5 causal variants, and found 1 risk-variant differing in the promoter activity between ER+ and ER-cell lines [21]. Therefore, there is a need for strategies to evaluate ER signaling in genetic backgrounds that reflect the diversity of women.

Models of ERα signaling in normal breast include *ex-vivo* culture of explants, organoid/3D culture, primary culture of mature luminal cells and ectopic expression of ERα. *Ex-vivo* cultures such as breast explant and tissue microstructures have been successful in retaining steroid hormone following tissue culture for up to 2 months [14,22–27]. In these cultures, normal tissue is used which retain breast cell types, tissue morphology, ERα expression, and response to hormone treatment. Recent studies have identified protocols for development of 3D breast organoids from primary HBECs and cancerous breast tissue [28–32]. Breast organoids contain luminal and basal epithelial cell populations, can organize into ductal structures, retain ERα expression, and are responsive to hormone treatment. However, propagation of breast organoids from normal breast tissue is limited as they can only be maintained for a short number of passages. For the same reason, freshly isolated primary luminal cells cells is limited by the amount of tissue obtained from donors and therefore only a finite number of experiments can be performed. HBEC models are reflective of the normal breast epithelium, but these are limited by senescence and loss of ERα in 2D culture. Multiple methods to bypass HBEC senescence were developed including transformation with HPV E6 and E7, human telomerase (TERT), Myc, SV40 large T-antigen, zinc-finger protein ZNF217, and BMI-1 [34–42]. However, the ERα expressing population in the HBECs rapidly decreased [43–45] in 2D culture possibly due to reduced adherence [46,47], faster growth of basal epithelial cells [47,43], stress responses in luminal epithelial cells [45], changes to the microenvironment [48], or epigenetic reprogramming that results in the loss of the differentiated phenotype [47,49]. Hence, there are currently few models of normal breast epithelial cells to study effects of ERα signaling in vitro and the variation that exists among women.

The goal of this study was to develop *in vitro* models for study of estrogen signaling using inducible expression of ERα in HBEC lines and utilize this model to study factors which may enhance or restrict signaling and sensitivity to estrogen. We used 4 stably immortalized HBEC lines: 76N TERT [50,51], MCF10A [52], HME-CC [53], and ME16C2 [53]. The 76N TERT, HME-CC, and ME16C2 HBEC lines were immortalized by expression of telomerase (*TERT*). The MCF10A line is spontaneously immortalized, likely through the loss of *p16INK4A*, *p14ARF*, and *p15INK4B* tumor suppressors [54]. All 4 HBEC lines are considered basal epithelial cell lines with expression of *KRT5* and *KRT14* [55], although some studies showed expression of luminal epithelial markers *KRT8* and *KRT18* in 76N TERT and MCF10A cells [56,57].

We developed a doxycycline inducible *ESR1* expression construct using a lentiviral backbone. In these HBEC lines with inducible expression of *ESR1*, the gene encoding ERα, we demonstrated ERα-dependent transactivation of target genes and cell proliferation. Conditioned media was not sufficient to elicit proliferation suggesting that proliferation was due to signaling intrinsic to the ERα-expressing cells. Transcriptomic analysis demonstrates a set of genes representing a core response to ERα signaling but also revealed variation in transcriptional patterns and estrogen-stimulated DNA double strand breaks among individuals. The ERα-mediated DNA damage was amplified in MCF10A-*ESR1* with heterozygous mutation of *BRCA1*^185delAG^.

## METHODS

### Cell culture

76N TERT, MCF10A, ME16C2, and HME-CC cells were obtained from Vimla Band, American Tissue Type Collection, Charles Perou, Christopher Counter, respectively, and have been described previously [58,59]. MCF10A cells heterozygous for the *BRCA1*^185delAG^ risk variant (NM_007294.4(BRCA1):c.68_69del) causing a frame-shift (p.Glu23fs) was obtained from Ben Park [60]. Cells were grown in F-media: DMEM-pyruvate (Gibco #11965-092), Ham’s F12 (Gibco #11765-054), 5% FBS (Omega scientific #FB-11), 250ng/mL hydrocortisone (Sigma #H4001), 10ng/mL human epidermal growth factor (Tonobo Biosciences #21-8356-U100), 8.6 ng/mL cholera toxin (Millipore Sigma #227035), 10 ug/mL human insulin (Sigma #I9278-5ML), and 1X antibiotic/antimycotic (Caisson Labs #ABL02-100ML). HEK239T cells were cultured in HEK293 growth media: DMEM:F12 (Sigma #D8900) with 10% FBS, 15ug/mL gentamycin (Gibco #15750-060), and 1X antibiotic/antimycotic. T47D cells were grown in DMEM:F12 supplemented with 10% FBS, 2mM L-Glutamine (Hyclone #SH30034.01), and 1X antibiotic/antimycotic. All cells were incubated at 37°C with 5% CO_2_ and passaged every 2-3 days.

### Generation of inducible HBEC-*ESR1* cell lines

Inducible *ESR1* construct (pIND-*ESR1*) was generated in our lab as described previously [3] using the pINDUCER14 vector modified to express the coding region of human gene encoding the ERα protein *ESR1* [61]. To generate lentiviral particles, HEK293T cells were grown overnight at 2.5×10^6^ cells per 60mm dish and transfected next day using 3.5µg pIND-*ESR1*, 3µg psPAX2 (Addgene #12260, gag, pol, and rev packaging vector), and 2µg pMD2.G (Addgene #12259, vsv-g packaging vector) in antibiotic free media with Lipofectamine 2000 (Thermo Fisher Scientific #11668019). After 24 hours, media was refreshed and replaced with HEK293T growth media. Media collected at 48 and 54h post-infection was filtered using a 0.45-micron filter (Corning #431220), mixed with F-media at 1:1 ratio and fed to HBECs for 24h. Proportion of GFP+ cells were estimated to determine the percentage of transfection. FACS was performed using pooled cells from each HBEC line in 1% FBS by selecting for GFP positive cells using FACSAria II (Becton-Dickinson). These cells are termed HBEC-*ESR1* cell lines. Uninfected parental HBECs were used as a negative control to set background fluorescence.

### Cell treatments for E2 response

Cells expressing ERα (HBEC-*ESR1* and T47D) were treated with clearing media consisting of MEM (Gibco #51200-038), 5% charcoal-dextran stripped serum (Omega Scientific #FB-04), and 2mM L-glutamine for 24 – 48h as noted. ERα expression was induced with 100ng/mL doxycycline (Sigma-Aldrich #D9891-1G). Treatments included either 10nM E2 (Sigma-Aldrich #E2758) or ICI 182 870 (Tocris #1047)) at 10nM or 1µM concentration from 10mM stock prepared in ethanol.

### ERE-luciferase Assays

HBEC-*ESR1* cells were plated in 24 well tissue culture plates at 1 x 10^5^ cells/well in F-media. Following overnight incubation, growth media was replaced with 0.3 mL/well Opti-MEM reduced serum media (Gibco #11058-021). Plasmid DNA and Lipofectamine 2000 reagent were diluted in Opti-MEM media before adding 200 µL of the mix to each well. HBEC-*ESR1* cells were infected with 3X ERE-TATA-Luc (Addgene, #11354) and Renilla using Lipofectamine 2000. After 6 hours of transfection, transfection media was replaced with 0.5 mL of clearing media. Next day, treatments of 10nM E2 or 10nM ICI with 0-200ng/mL doxycycline was added. Promega Dual Luciferase Reporter Assay (Promega # E1910) was used to perform luciferase assays. Cells were lysed in 1x Passive Lysis Buffer after 24-hour treatment and lysates stored at –20°C until reading. Luciferase and Renilla activity in lysates were determined by using the Polar Star Optima plate reader (BMG Labtech). Luciferase values for each well was normalized to the amount of Renilla activity.

### Western blot

HBEC-*ESR1* cells and MCF7 were treated as described above and lysed in ice cold RIPA lysis buffer [50 mM Tris–HCl pH 8.0, 150 mM NaCl, 1 mM EDTA, 1% Triton X-100, 1% Sodium deoxycholate, 0.1% SDS, 1% phosphatase inhibitor #3 (Sigma-Aldrich #P0044)]. Lysates were centrifuged at 13,000 rpm for 15 minutes at 4°C to remove cellular debris. BCA protein assay (Thermo Scientific #23225) was performed to quantify protein concentration. For each cell line, equal amounts of protein (28 µg) were separated using SDS-PAGE on 10% acrylamide gels under denaturing conditions and blotted onto PVDF membrane (Millipore #IPVH00010). Blocking was performed using 5% nonfat dry milk in TBST (10mM Tris-HCl pH 7.5, 150mM NaCl, 0.05% tween-20) for 1 hour. Blots were incubated with 1:100 anti-ERα (Abcam [Sp1] #ab16660, Lot #GR3202692-2) overnight at 4°C.

The next day, blots were washed 3 times with TBST and then incubated for 1 hour with HRP-conjugated secondary antibody (1:5000, GE Healthcare #NA934V). Protein bands were detected using enhanced chemiluminescence solution and imaged on G-box (Syngene). The blots were washed again with TBST, incubated with anti-β actin (1:5000, Sigma #A1978) overnight at 4°C, re-washed and incubated with secondary antibody(1:5000, GE Healthcare #NA931C), and detection were performed as described above. Expected molecular weights were 67 kDa for ERα and 42 kDa for β actin.

### Cell proliferation assay

HBEC-*ESR1* cells were plated in clearing media on five 96-well plates at the cell density of 10000 cells per well. The next day, cells were treated with 100ng/mL doxycycline and vehicle control (Dox+Control, 10nM E2 (Dox+E2) or 10nM ICI (Dox+ICI). Each day, Alamar blue reagent was added (final concentration 10%) and incubated for 4 hours. Plates were read at 570nm and 600nm in a BioTek Synergy 2 plate reader (BioTek). Media was refreshed on day 3 on remaining plates. Plates were read for 4 days. Percent Alamar blue reduction was calculated using the Alamar blue protocol (Invitrogen # DAL1100).

### Conditioned growth media

Cells were plated at specified density (HBEC-*ESR1* cells at 1.6×10^6^ & T47D cells at 1.2×10^6^ cells per T75 flask) in clearing media (10% CSS, 2mM L-Glutamine, 10ug/mL insulin, and MEM media) for 48-72 hours. 10ug/mL insulin, and MEM media. 48 hours after treatment, media was collected and replaced with fresh media. Conditioned media was filtered with a 0.2-µM syringe filter, diluted 1:1 in fresh media, and used to treat cells in proliferation assays. After an additional 48 hours, this process was repeated, and conditioning cells were discarded.

### Immunofluorescence

HBEC-*ESR1* cells were plated in 8-chambered slides (CELLTREAT #229168) at 30,000 cells per chamber in F media. After 24h of growth, F-media was replaced with phenol red free F-media with 10% CSS (PRF-CSS) along with 100ng/mL of doxycycline. After 24h of PRF-CSS treatment, cells were treated with either Dox+Control or Dox+E2 in PRF-CSS media for 20h. If required, Mirin (50µM), NU4771 (10µM) or Vehicle Control was added and incubated an additional 4h. Cells were treated with 3.7% paraformaldehyde (Electron Microscopy Sciences #15710-S) for 30min, washed with PBS twice and stored at 4C till further processing. For immunostaining, the fixed 8-chambered slides were quenched by incubating the slides in 0.1N Glycine at pH 8 for 20min followed by 2 washed in PBS. Then the slides were incubated in a blocking solution of 10% Donkey serum (Sigma-Aldrich #D9663-10ML) with 0.3% Trition-X-100 in PBS pH 8 for 1 hour. The slides were incubated in primary antibodies CyclinA2 (Thermo Firsher Scientific # MA1154) and/or γH2AX (Cell Signalling # 9718S) for 2h at RT, followed by 3 washes in PBS, 1 hour incubation with anti-Rabbit Alexa-Fluor 594 (Thermo Fisher Scientific # A32754) and anti-Mouse Alexa-Fluor 647 (Thermo Fisher Scientific #A32787) antibodies and 3 washes with PBS. Coverslips were mounted on the slides with mounting media (Vectaschild Vibrance with DAPI #H-1800). The slides were imaged in a NIKON A1 Spectral detector confocal microscope at 60x. Image analysis was performed with NIKON AR analysis software with GA3 module. Nuclei were outlined with DAPI mask and γH2AX and CyclinA2 foci were quantified within nuclear area.

### RT-qPCR

RNA isolation was performed using TRIzol according to manufacturer’s recommendation (ThermoFisher synthesis kit (New England Biolabs #E6560L). RT-qPCR was performed in a thermocycler (CFX96 Real-Time thermocycler, BioRad). See **Supplementary Table 1** for primer sequences. Expression is relative to an inter-run calibrator (IRC) consisting of pooled cDNA from a subset of human breast tissue samples.

### RNA-Seq analysis

RNA was sent to LC Sciences (Houston, TX) to perform sample QC, library preparation, and poly(A) RNA-sequencing (150 bp PE, 40 million reads). At LC Sciences, The total RNA quality and quantity were analysis of Bioanalyzer 2100 and RNA 6000 Nano LabChip Kit (Agilent, CA, USA) with RIN number >7.0. Approximately 10 ug of total RNA was subjected to isolate Poly (A) mRNA with poly-T oligo attached magnetic beads (Invitrogen). Following purification, the poly(A) mRNA fractions is fragmented into small pieces using divalent cations under elevated temperature. Then the cleaved RNA fragments were reverse-transcribed to create the final cDNA library in accordance with a strand-specific library preparation by dUTP method. The average insert size for the paired-end libraries was 300±50 bp. And then we performed the paired-end 2×150bp sequencing on an Illumina Hiseq 4000 at LC Sciences following the vendor’s recommended protocol.

RNA-Seq data was verified using md5 to ensure data integrity. Read quality was assessed with FASTQC (v0.11.5) and compiled using MultiQC (v1.4)[62]. Reference genome was indexed with STAR (v2.7e) [63] from GRCh38/hg38 using human GENCODE gene set release 39 using RSEM[64]. Genomic alignment and read quantification were done with rsem-calculate-expression. RSEM output as gene level result files were imported to R using tximport (v1.22.0). Genes with low read counts (< 5) in more than 3 samples were filtered out. Differential gene expression analysis was performed using DESeq2 (v1.34.0) [65] using pair-wise comparison. Differentially expressed gene list was obtained using an adjusted p-value cutoff of 0.05.

Gene Set Enrichment Analysis was performed using GSEA (v4.3.2)[66,67]. Expression matrix of Log2 count from the Dox-E2 and Dox+Control treated HBEC-*ESR1* and the Hallmark gene-sets [68] were used for MSigDB gene sets were used with an FRD q-value of 0.25.

## RESULTS

### Generation of HBEC-*ESR1* lines using pIND-*ESR1*

We generated 4 HBEC lines with inducible *ESR1* to obtain controlled expression of the ERα protein. A modification of the pINDUCER14 vector was used to express the human *ESR1* sequence coding for the ERα protein [3,61]. This construct (pIND-*ESR1*), contains an EF1α promoter which drives constitutive expression of the reverse tetracycline-transactivator (rtTA) and eGFP. The *ESR1* coding sequence with an N-terminal FLAG tag is preceded by a tetracycline response element (TRE2) (**Figure 1A**). Each HBEC cell line was infected with pIND-*ESR1* and the cells were enriched for GFP+ cells by FACS (**Supplementary** Figures 1-4). Following FACS, the cell lines were designated 76N TERT-*ESR1*, MCF10A-*ESR1*, ME16C2-*ESR1*, and HME-CC-*ESR1*. The 76N TERT-*ESR1* were 90% GFP+, MCF10A-*ESR1* were 70% GFP+, HME-CC-*ESR1* were 90% GFP+, and ME16C2-*ESR1* were 98% GFP+ (**Supplemental Figure 5**).

**Figure 1:**
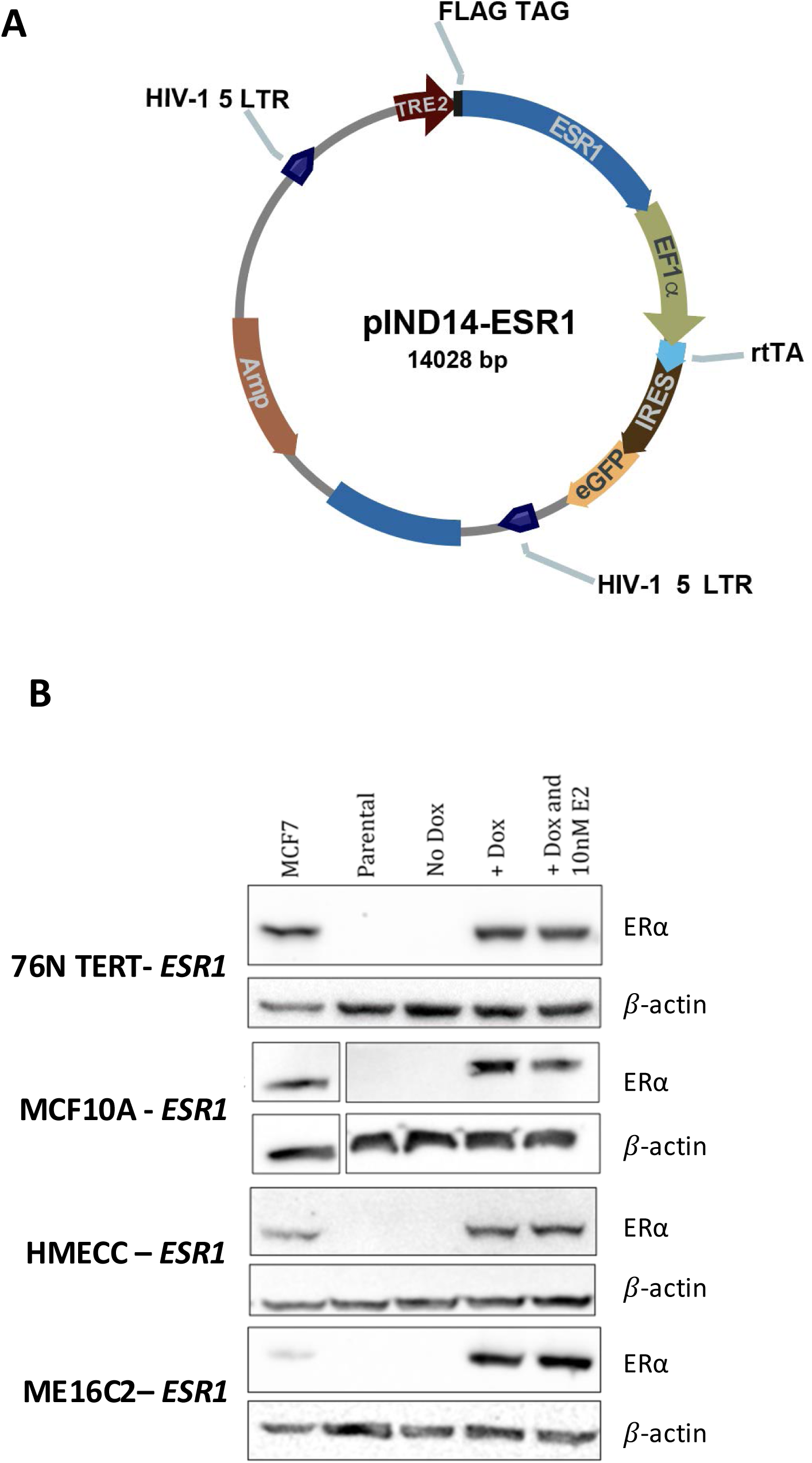
Generation of HBECs expressing inducible ERα (HBEC-*ESR1*). **A**) Vector map of pIND-*ESR1*. rtTA and GFP are constitutively expressed by EF1α promoter and *ESR1* is expressed by TRE2 promoter. **B)** ERα protein levels in 76N TERT-*ESR1*, MCF10A-*ESR1*, HMECC-*ESR1* and ME16C2-*ESR1*. GFP^+^ cells pooled and treated with 100ng/ml doxycycline show similar levels of ERα for each HBEC-*ESR1* as MCF7.

Expression of ERα was compared with MCF7 breast cancer cell line. The parental HBEC lines and un-induced HBEC-*ESR1* cells lacked expression of ERα. All 4 HBEC-*ESR1* showed ERα levels at 67 KDa with 100ng/mL doxycycline treatment (**Figure 1B**). ERα protein levels were normalized to β-actin. 10nM E2 treatment did not downregulate ERα expression in these cell lines (**Figure 1B**, **Supplemental Figures 6**). 76N TERT-*ESR1*, MCF10A-*ESR1*, and HME-CC-*ESR1* expressed ERα approximately equal to that in MCF7 cells. The ME16C2-*ESR1* line expressed ERα at 6-fold more than MCF7 cells. These results indicated that all 4 HBEC-*ESR1* cell lines had minimal background levels and express ERα proteins at levels sufficient to elicit E2-induced responses.

### E2-induced transcriptional activation in HBEC-*ESR1* cells

Luciferase assays were performed to evaluate the activity of ERα signaling with increasing concentrations of doxycycline. Induction of *ESR1* expression with doxycycline (Dox) together with addition of 10nM E2 stimulated large increases in transactivation of the ERE-luciferase construct in all HBEC-*ESR1* cell lines demonstrating that responses were both receptor– and ligand-dependent (**Figure 2A-D**). In the absence of doxycycline, E2 treatment failed to induce the luciferase reporter in 76N TERT-*ESR1* and HME-CC-*ESR1* cells and showed dose-dependent increases in transactivation with 100, 150 and 200ng/mL doxycycline consistent with the induction of ERα protein (**Figure 2A,C**). In MCF10A-*ESR1* cells, addition of E2 resulted in detectable increases in luciferase activity without Dox, suggesting there may be leaky expression of *ESR1* in these cells. There were progressive increases in luciferase expression with 100 and 150 ng/mL with a maximum at 200 ng/mL Dox **(Figure 2B)**. There was significant background activity in the ME16C2-*ESR1* cells treated with E2 in the absence of Dox. The ME16C2-*ESR1* cells had the highest level of ERα protein which appears to cause ligand-independent transactivation. The cells also had the highest fold increases in luciferase activity when treated with E2 reaching ∼37 and 60-fold increases with 100 and 150 ng/mL Dox **(Figure 2D)**. The level of ERα protein induced by 100 ng/mL of doxycycline was similar to that found in MCF7 cells for 76N TERT-*ESR1*, MCF10A-*ESR1* and HME-CC-*ESR1* cells, whereas ERα in ME16C2-*ESR1* cells was over 6-fold higher compared to MCF7 cells **(Figure 1B, Supplemental Figure 6)** and contribute to the greater activity of the luciferase reporter. Together these data show that 100 ng/mL of Dox was sufficient to induce ERα and elicit ligand-dependent transactivation of the reporter gene.

**Figure 2:**
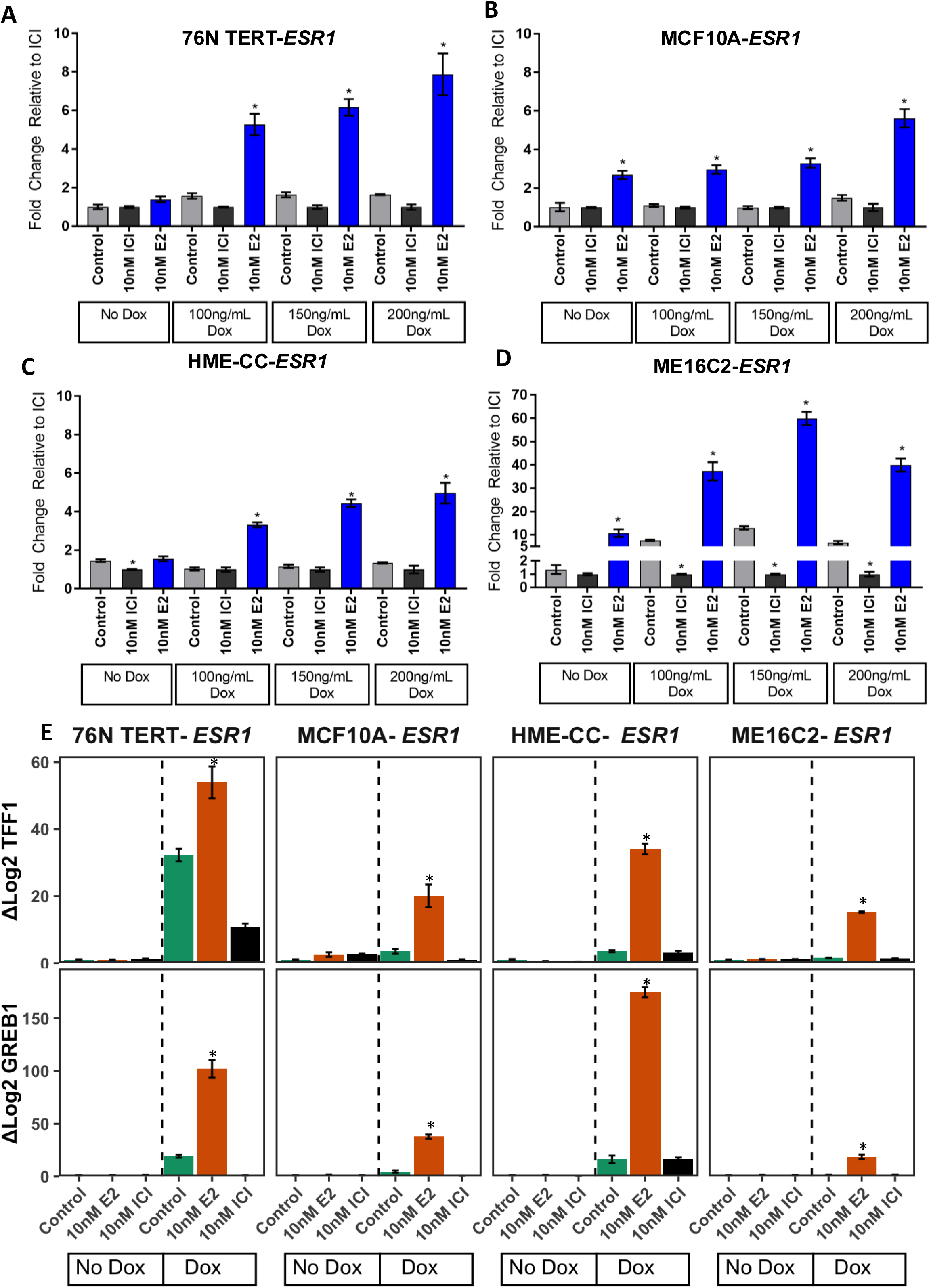
Transactivation of inducible ERα in all 4 HBEC-*ESR1* lines. ERα transactivation with 3X ERE-TATA-Luc luciferase reporter assays were performed in **A)** 76N TERT-*ESR1*, **B)** MCF10A-*ESR1*, **C**) HME-CC-*ESR1*, and **D)** ME16C2-*ESR1* cell lines. Control response (grey bar) was compared with E2 response (blue bar) and ICI response (black bars). Without any ERα induction (No Dox) E2 responses are not significantly different from Control in 76N TERT-*ESR1* **(A)** and HME-CC-*ESR1* **(C)** but significantly increased in MCF10A-*ESR1* (**B**) and ME16C2-*ESR1* **(D)**. Increase in doxycycline levels led to higher transactivation in all cells. ICI response was similar to Control in all levels of doxycycline. (\**p* < 0.05). **E)** Expression of *TFF1* and *GREB1* are ERα inducible in HBEC-*ESR1* lines. In No Dox condition, no expression of *TFF1* and *GREB1* was detected in qRT-PCR. 76N TERT-*ESR1* and HME-CC-*ESR1* expressed *TFF1* and *GREB1* with Dox+Control. All 4 cell lines showed strong induction of *TFF1*and *GREB1* following with Dox+E2 treatment. Conversely, Dox+ICI treatment repressed both gene expression.

Transactivation of ERα-target genes within cells require ERα-dependent recruitment of transcriptional coactivators to estrogen-response elements in promoters and are sustained through 24h [69–71]. Therefore, we examined E2 induction of the *TFF1* and *GREB1* cellular genes following treatment with Dox+E2 (**Figure 2E**). In the absence of Dox, E2 failed to induce expression of either *TFF1* or *GREB1* in any cell line. In contrast, E2+Dox stimulated strong induction of *TFF1* and *GREB1* demonstrating ligand-dependent transcriptional activation of these cellular genes. While all 4 HBEC-*ESR1* cell lines were responsive to E2, the sensitivity differed among the lines and genes. The 76N TERT-*ESR1* cells had the largest fold induction of *TFF1* and *GREB1* expression with E2+Dox but also had significant expression with Dox alone compared to the No Dox samples. In contrast, the MCF10A-*ESR1* and HME-CC-*ESR1* cells had strong increases in *TFF1* and *GREB1* expression with E2+ Dox with only modest background when ERα expression was induced by Dox in the absence of the E2 ligand. Although ME16C2-*ESR1* cells had the highest levels ERα that were ∼6-fold greater than levels found in MCF7 cells (**Supplemental Figure 6**) and extremely high transactivation of the ERE-luciferase reporter, it elicited weaker transactivation of the cellular *TFF1* and *GREB1* genes when treated with E2+ Dox (**Figure 2E**). While unexpected, the abundance of ERα protein may have limited the availability of coactivators to form complexes necessary to bind to promoters of cellular genes. Alternatively, responses may reflect differences in genetic polymorphisms and chromatin resulting in private patterns of ERα binding and transactivation among individuals.

### HBEC-*ESR1* cells show proliferative response to E2 that is not dependent on secreted factors

ERα transcriptional activation leads to increased proliferation of ERα+ breast cancer cell lines [72,73]. Similarly, induction of ERα by Dox+E2 increased proliferation of 76N TERT-*ESR1*, MCF10A-*ESR1*, and HME-CC-*ESR1* lines (**Figure 3A-C**). Proliferation was lower in the absence of E2 despite expression of ERα (Dox+Control) indicating ligand-dependence. ME16C2-*ESR1* cells did not show significant difference in proliferation with Dox+E2 over Dox+Control (**Figure 3D**). Although ERα signaling was reconstructed in all 4 HBEC-*ESR1* cell lines based on expression of *TFF1* and *GREB1*, proliferative responses were detected in only 3 cell lines which could be due to the failure to transcriptionally activate specific growth factors.

**Figure 3:**
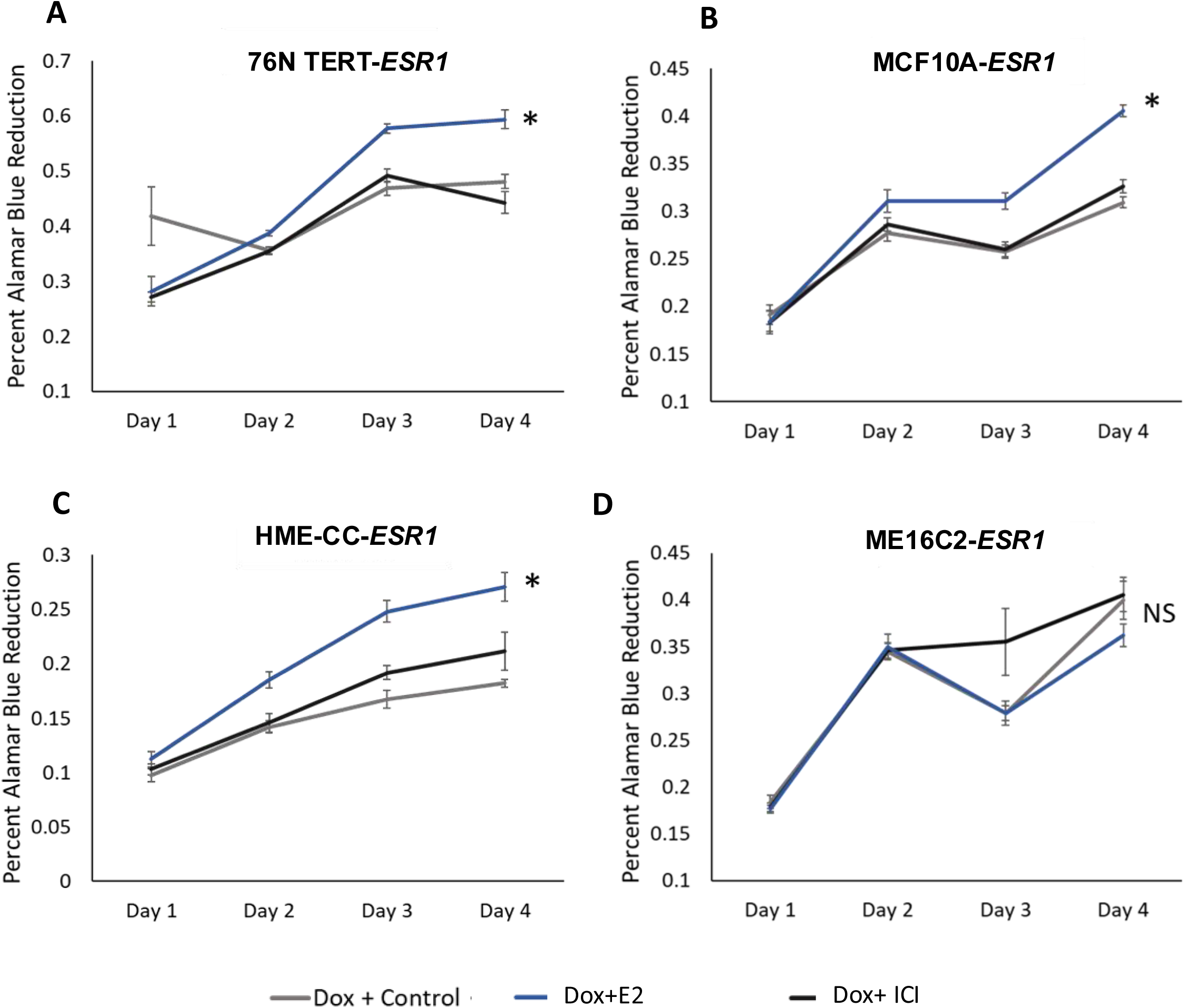
E2-induced proliferation in HBEC-*ESR1*. Cell proliferation of HBEC-*ESR1* after treatment with Dox+Control, Dox+E2 or Dox+ICI. **A**) 76N TERT-*ESR1*, **B**) MCF10A-*ESR1*, **C**) HME-CC-*ESR1* show increase in proliferation with Dox+E2. Dox+ICI shows same level of proliferation as Dox+Control. **D**) ME16C2-*ESR1* showed no difference in cell proliferation between treatments. (* p < 0.05). Error bars indicate SEM.

Proliferation of breast epithelial cells during normal tissue development is mediated by paracrine secretion of factors from hormone-sensing (ERα+) cells, but in breast cancer cells, E2 can stimulate secretion of autocrine growth factors. To determine if E2-induced proliferation observed in HBEC-*ESR1* lines was due to cell intrinsic factors or production of secreted growth factors, we tested the effect of conditioned media from cells (**Figure 4A**). Cells responsive to E2-induced proliferation (76N TERT-*ESR1* and MCF10A-*ESR1*) were used to produce conditioned media (CM). Parental cell lines without ERα (76N TERT, MCF10A) were then placed in the CM and proliferation was measured. CM was collected from donor 76N TERT-*ESR1* and MCF10A-*ESR1* lines after treatment with Dox+Control or with Dox+E2 to induce ERα. CM from cells treated with Dox+ICI was used to block ERα signaling. Although 76N TERT-*ESR1* and MCF10A-*ESR1* cells proliferate in response to Dox+E2, CM from these cells failed to increase proliferation of the parental cells lacking ERα and did not differ from the Dox+Control or Dox+ICI treatments (**Figure 4B&C**). CM from the MCF10A-*ESR1* donor cells treated with Dox+Control or Dox+ICI treatments also failed to increase proliferation in the ERα+ T47D breast cancer cells (**Figure 4D)**. Proliferation of T47D cells was observed in CM from the donor cells when E2 was present (Dox+E2) demonstrating that the added E2 was required in these cells. These data demonstrate that the HBEC-*ESR1* cells do not stimulate secreted growth factors. Therefore, inducible expression of ERα was sufficient to mediate E2-stimulated proliferation in 76N TERT-*ESR1*, MCF10A-*ESR1* and HME-CC-*ESR1* cells through intracellular ERα signaling.

**Figure 4:**
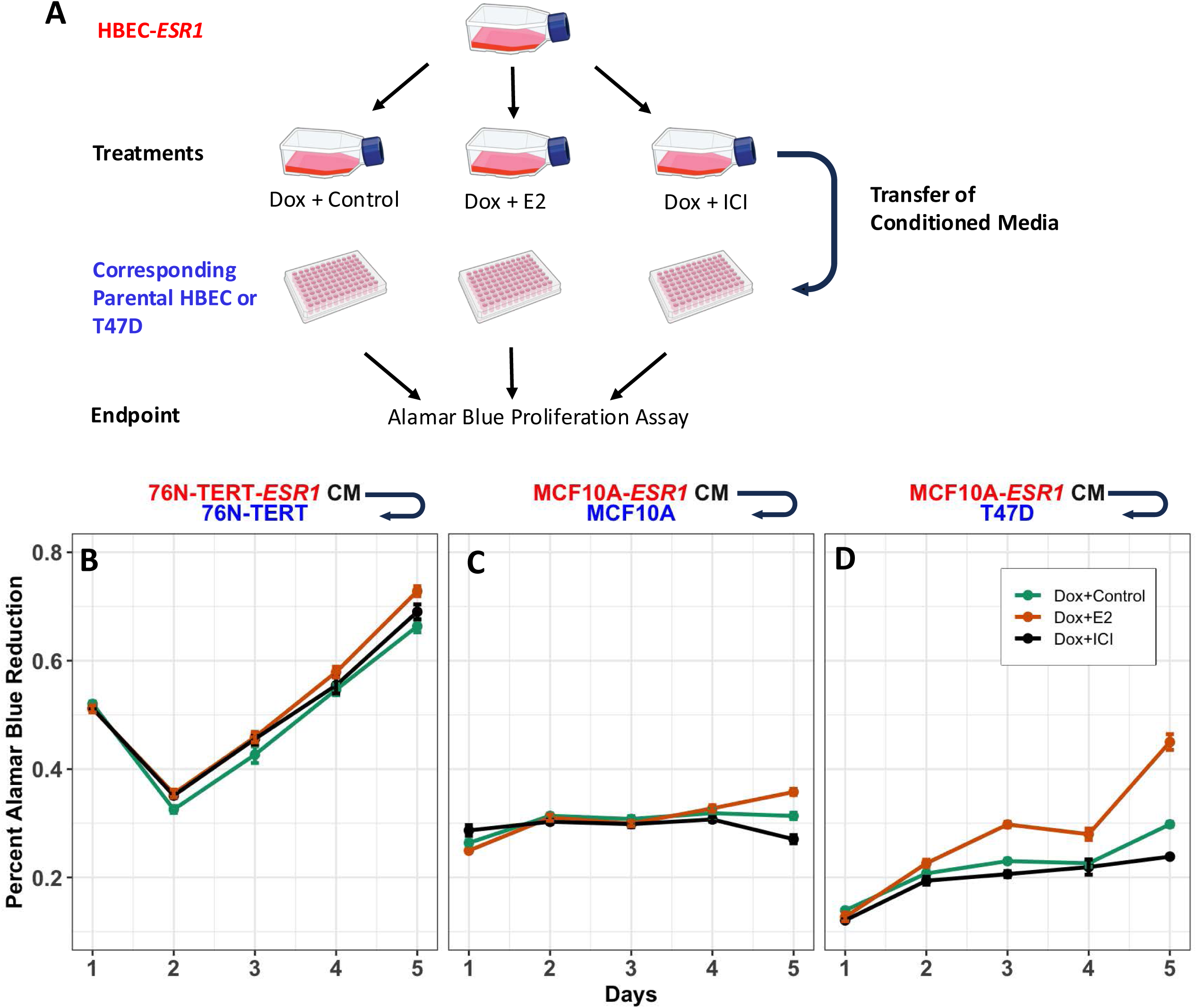
Cell intrinsic factors contribute to E2 mediated cell proliferation. **A**) Experimental design for conditioned growth experiment. Donor HBEC-*ESR1* (i.e. either 76N TERT-*ESR1* or MCF10A-*ESR1*) were treated with Dox+Control, Dox+E2 or Dox +ICI. Conditioned media (CM) from these cells were collected at 48h and 72h, filtered, diluted 1:1 with minimal growth media and added to the receiver cell as shown in **B-D**. Proliferation of the receiver cells were monitored for 5 days with Alamar blue reduction. **B**) Proliferation of parental 76N TERT cell line with conditioned media from 76N TERT-*ESR1* cells. **C**) Proliferation of parental MCF10A cell line with conditioned media from MCF10A-*ESR1* cells. **D**) Proliferation of T47D cells treated with conditioned media from MCF10A-*ESR1* cells.

### E2 mediated transcriptional response

Although expression of ERα was sufficient to elicit E2-dependent transcriptional responses in target genes and proliferation, the responses varied among the HBEC-*ESR1* lines. RNA-Seq was used to study differences in the E2-responsive transcriptomes among the HBEC-*ESR1* lines. Transcriptional responses to E2 at 24h was used to enrich for direct targets of ERα. The top 1000 genes that were highly variable among the 4 HBEC-*ESR1* cell lines and 3 treatments were used for principal component analysis. The 2^nd^ to 3^rd^ principal components distinguished the patterns of transcription that are unique to each cell line (**Figure 5A**). The effect of treatments on gene expression patterns was observed using 4^th^ and 5^th^ principal components, which clusters the treatment groups (**Figure 5B**). The variation within treatments suggests that transcriptional patterns for the estrogen-responsive genes vary among the individual donors from whom the HBECs were derived. Adjusting for baseline differences between the cell lines, we performed a paired design of differential gene expression analysis to compare between Dox+Control and Dox+E2. A total of 682 genes were found to be differentially regulated with an adjusted p-value cut-off of p=0.05, with 418 upregulated and 264 downregulated **(Figure 5C)**. 43% of the HBEC-*ESR1* estrogen-induced differentially expressed genes (DEG) overlapped with either MCF7 and/or T47D (**Figure 5D**). Significant inter-individual variation in E2-responsive genes is also detected between the MCF7 and T47D breast cancer cell lines as well as among the HBEC-*ESR1* cell lines. However, a core set of E2-responsive genes showed consistent responses across the HBEC-*ESR1* lines and inverse responses were observed in the ICI treatment confirming the ERα-dependent expression (**Figure 5E**). This core set of 682 genes showed a similar expression pattern in MCF7 and T47D breast cancer cell lines. Gene set enrichment analysis (GSEA) identified 8 gene-sets which were positively associated and 1 negatively associated with Dox+E2 treatment. The Hallmark Estrogen responses were the top 2 gene sets confirming the enrichment of ERα target genes and pathways. Additionally, GSEA identified pathways contributing to cell division such as E2F Targets, G2M checkpoints, *Myc* Targets, Oxidative phosphorylation, Spermatogenesis and DNA repair (**Figure 5F**). Among the DNA Repair genes, enrichment scores were greatest for genes that were induced when ERα signaling was engaged **(Figure 5G)**. Therefore, inducible expression of ERα in HBEC lines recovers core transcriptional responses that are shared among individuals and ERα+ breast cancer cell lines.

**Figure 5.**
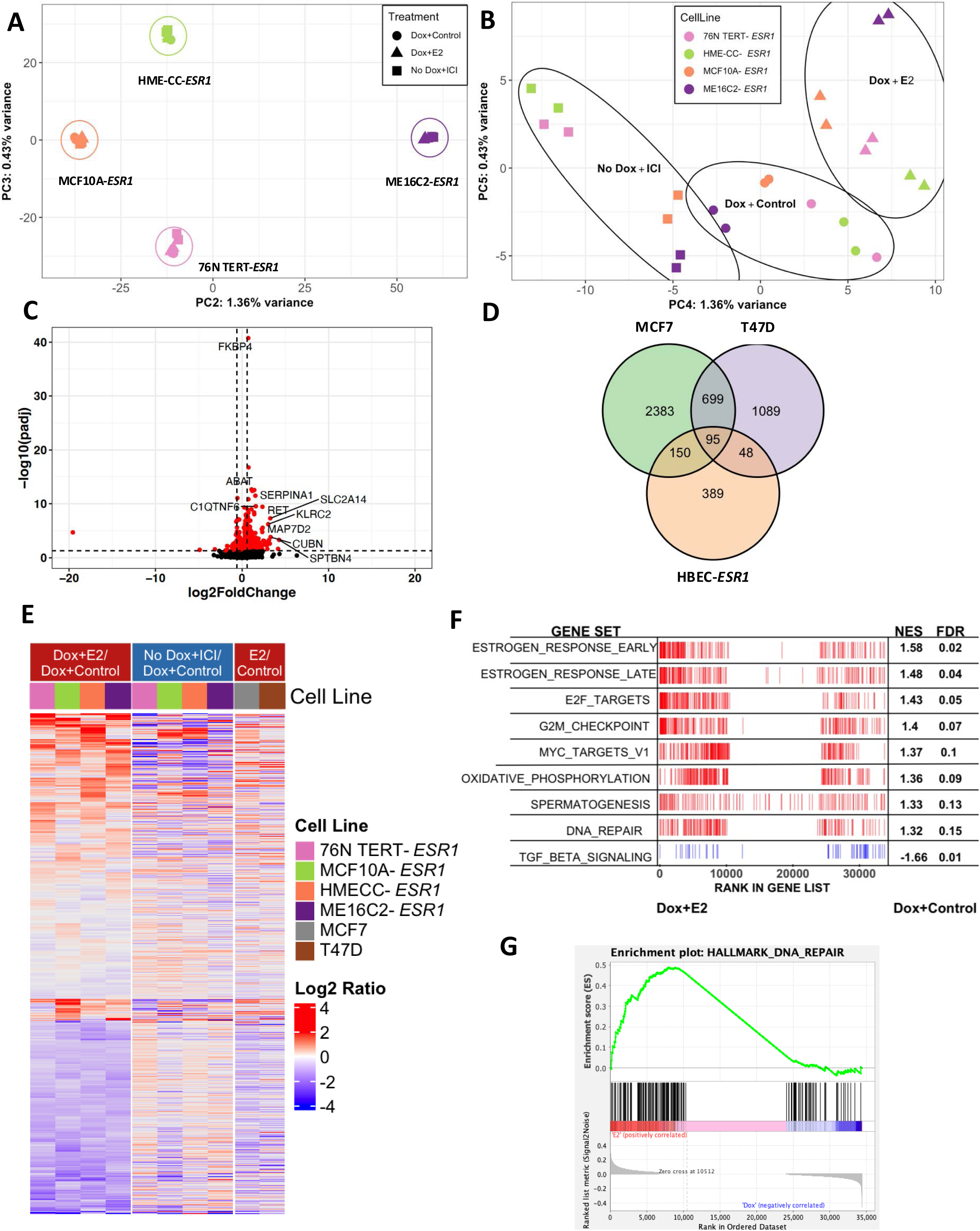
Transcriptional analysis of E2 response in HBEC-*ESR1* cells. **A)** PCA of top 1000 variant genes of estrogen treatment (Dox+E2) vs vehicle control (Dox+Control) with all HBEC-*ESR1* show 682 genes which are differentially expressed (red circle) (FRD < 0.05). Top 10 differentially expressed genes (DGE) are labelled with Gene symbols. **D)** Overlap of HBEC-*ESR1* estrogen response DGEs with MCF7 and T47D. **E)** Heatmap of 682 HBEC-*ESR1* DEGs showing inverse trend in E2 and ICI treatments, and similar trend in BC cell lines (MCF7 and T47D). **F)** GSEA analysis of HBEC-*ESR1* Dox+E2 vs Dox+Control expression data identified 8 pathways positively enriched and 1 negatively enriched in Dox+E2 treated cells (FDR 0.25). **G)** Enrichment plot of DNA repair pathway comparing Dox+E2 vs Dox vs Dox+Control.

### ERα dependent E2 mediated DNA damage

Upregulation of DNA repair pathways in RNA-Seq analysis in response to E2 treatment suggests ongoing DNA damage. We next studied DNA double strand breaks (DSBs) following induction of ERα and treatment with E2 for 24 hours. Median γH2AX foci per nucleus was used as a measure DSBs. Dox+E2 treated MCF10A-*ESR1*, 76N TERT-*ESR1* and ME16C2-*ESR1* cells showed significantly higher median DNA damage when compared with their respective Dox+Control treated cells (**Figure 6A&B**). HME-CC-*ESR1* cells showed higher median γH2AX foci in both Dox+Control and Dox+E2 treatments compared to the other 3 HBEC-*ESR1* cell lines. However, there was not a significant increase in DSB with Dox+E2 treated in HME-CC-*ESR1* cells compared to the Dox+Control. To determine the relationship between DNA damage due to replication, MCF10A-*ESR1* were stained for CyclinA2 to distinguish cells in G1 and S/G2 phases. At 24h after treatment with Dox+E2, MCF10A-*ESR1* cells showed dramatically higher γH2AX foci in S/G2 cells (CylinA2^high^) compared to Dox+Control cells. Nuclei of the S/G2 phase cells showed the highest levels of DSB (∼10 fold over Dox+Control) (**Figure 6C&D**). However, the G1 cells (CylinA2^low^) also exhibited significantly higher median levels of DSBs. Therefore, ERα signaling induces measurable increases in DSB as well as stimulating replication-associated DNA damage.

**Figure 6.**
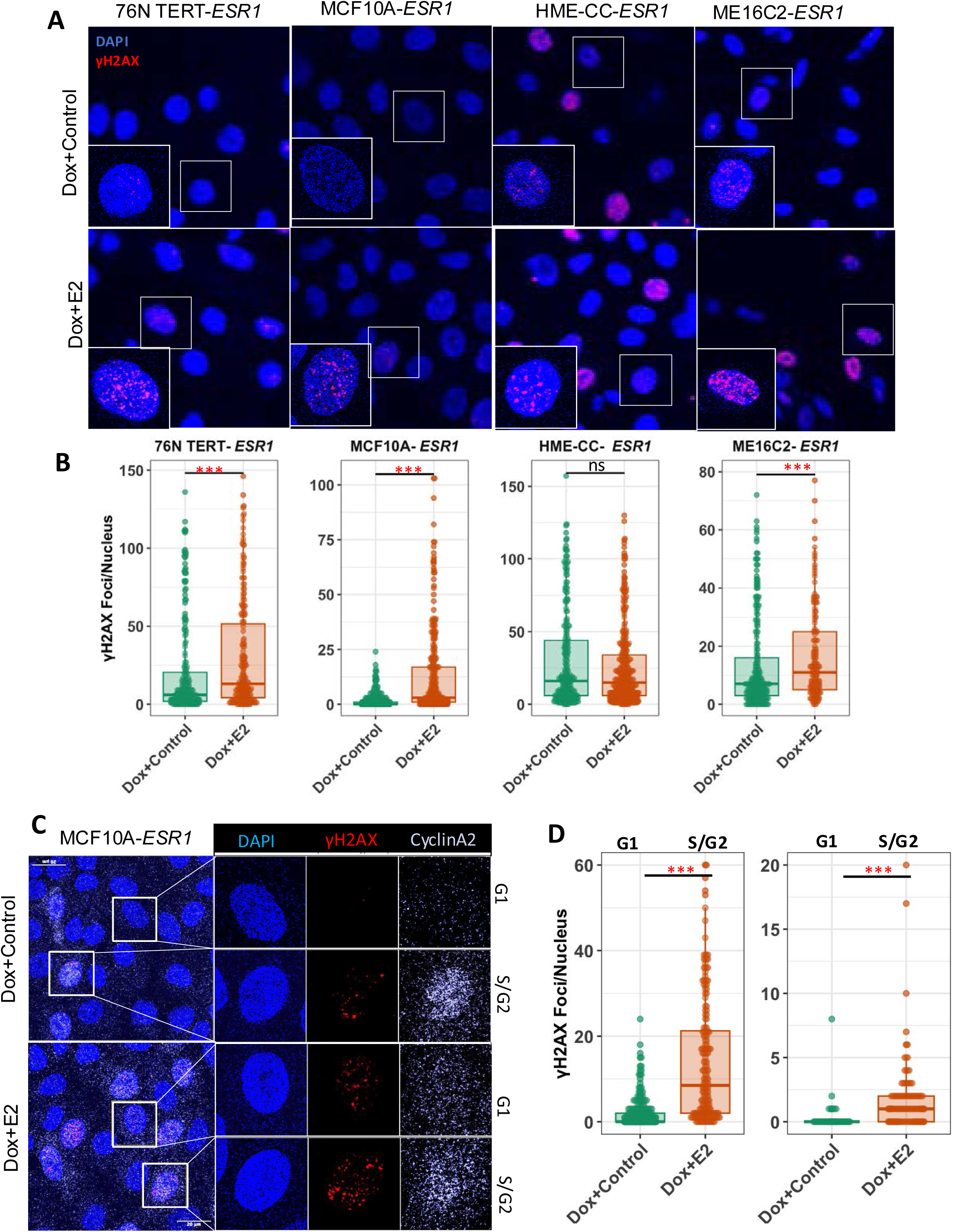
ERα-mediated DNA damage in HBEC-*ESR1*. **A)** Immunofluorescence staining of γH2AX (red) and nucleus (blue) in HBEC-*ESR1* cells treated with Dox+Control or Dox+E2. **B)** Quantification of nuclear γH2AX in **A**. 76N TERT-*ESR1*, MCF10A-*ESR1* and ME16C2-*ESR1* showed increased nuclear γH2AX foci Dox+E2 over Dox+Control. No increase in γH2AX was observed in HME-CC-*ESR1*. **C)** Immunofluorescence staining of nucleus(blue), γH2AX(red) and CyclinA2 (white) in MCF10A-*ESR1* showing DNA damage in CyclinA2^high^ (S/G2) and CyclinA2^low^ (G1)cells. **D**) Quantification of **C** showing increase in nuclear γH2AX in Dox+E2 over Dox+Control in both S/G2 and G1 cells. S/G2 cells show 10-fold higher DNA damage over G1 Dox+E2 treated cells.

Given that E2-mediated transactivation increases both DNA damage and DNA repair pathways, impairment of DSB repair pathways may potentiate E2-mediated DNA damage. HR requires MRE11 while NHEJ relies on DNA-PK. Therefore, inhibitors of MRE11 (Mirin) and DNA-PK (NU7441) were used to block HR and NHEJ, respectively. MCF10A-*ESR1* and ME16C2-*ESR1* cells were selected as they both showed increases in DSBs in response to Dox+E2 but differed in levels of ERα protein and proliferative responses. The cells were treated with Mirin or NU7441 for 4h following E2 treatment then stained for γH2AX (**Figure 7A)**. In MCF10A-*ESR1* cells, Mirin significantly increased the γH2AX foci/nucleus indicating a role for HR in resolving the ERα-mediated DSBs while inhibition of NHEJ with NU7441 had a more modest effect (**Figure 7B**). The higher levels of ERα in ME16C2-*ESR1* cells had higher baseline levels of γH2AX foci as observed previously and inhibition of either HR or NHEJ resulted in further increases in nuclear γH2AX foci (**Figure 7C**). Therefore, both HR and NHEJ play roles in mitigating the DSBs associated with ERα signaling.

**Figure 7.**
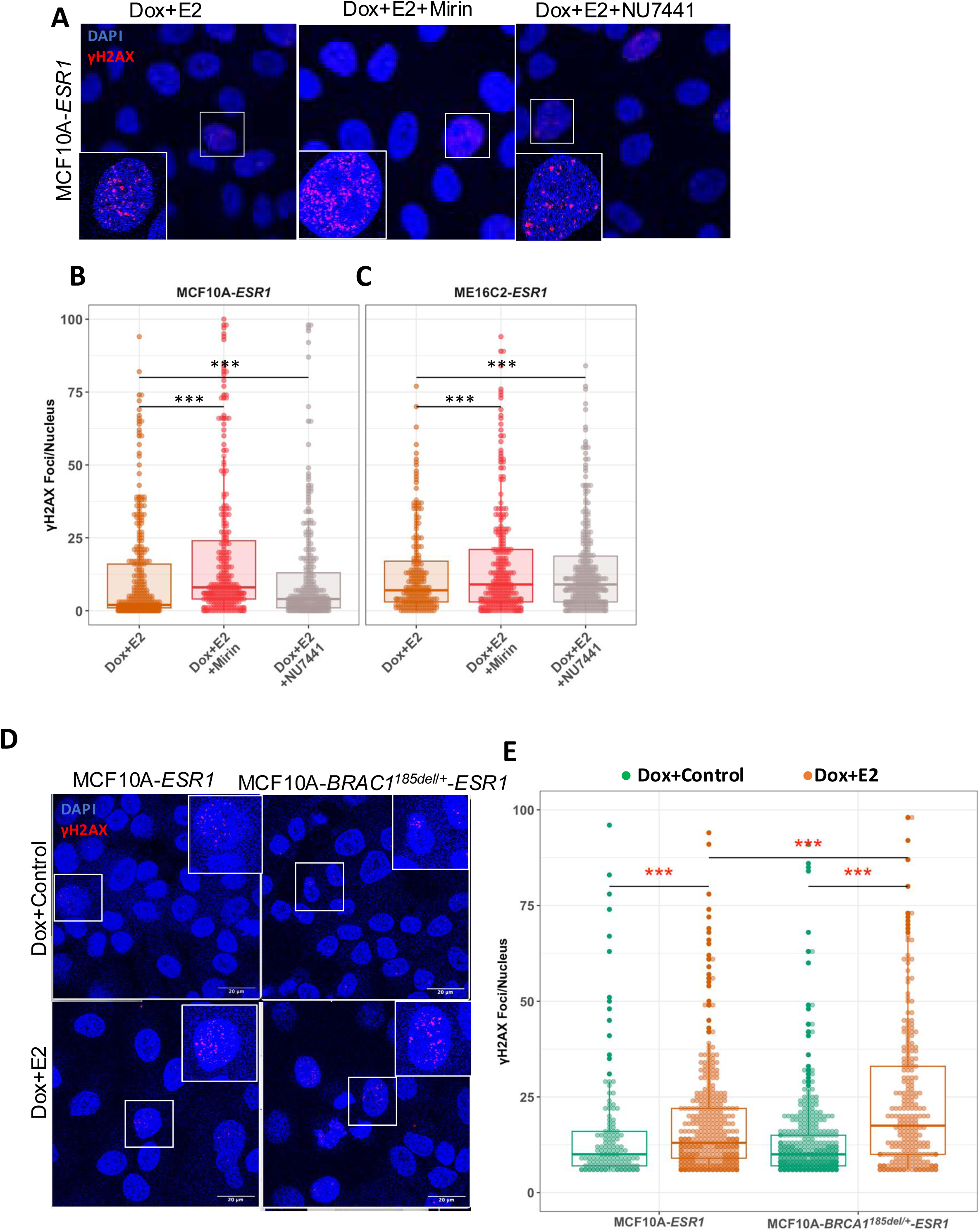
Inhibition of DNA repair pathway increases ERα-mediated DSB. **A**) Immunofluorescence staining of γH2AX(red) and nucleus(blue) MCF10A-*ESR1* cells showing DNA damage following treatments with 24h of E2 and 4h of DNA repair pathway inhibitors. **B**) Quantification of nuclear γH2AX in MCF10A-*ESR1* shows increase in DNA damage following MRE11 inhibitor Mirin and DNA-PKC inhibitor NU7441 treatment. **C**) Quantification of nuclear γH2AX in ME16C2-*ESR1* showing the similar levels of increase in DSB following treatment with Mirin or NU7441. **D**) Immunofluorescence staining of γH2AX(red) and nucleus(blue) of MCF10A-*ESR1* and MCF10A-*BRCA1*^+/185delAG^-*ESR1*. **E**) Quantification of **A** showing comparable levels of nuclear γH2AX foci in Dox+Control treated cells, but higher in Dox+E2 treated MCF10A-*BRCA1*^+/185delAG^-*ESR1* compared to Dox+E2 treated MCF10A-*ESR1*.

Inherited defects in DSB repair preferentially increases risk of breast cancer compared to other tissues. To examine the effect of haploinsufficiency in *BRCA1* on ERα-mediated DNA damage, MCF10A-*BRCA1^185del/+^* cells [60,74] were transduced with pIND-*ESR1*. The frequency of γH2AX foci in the isogenic MCF10A-*ESR1* and MCF10A-*BRCA1^185del/+^-ESR1* cells did not differ significantly following treatment with Dox+Control to induce ERα. Treatment with Dox+E2 to activate ERα signaling resulted in significant increases in γH2AX in both cell lines, but the magnitude of DSBs was greater in MCF10A-*BRCA1^185del/+^-ESR1* compared to MCF10A-*ESR1* (**Figure 7D,E**). Therefore, impairment of DSB repair pathway by inhibition of HR/NHEJ or haploinsufficiency in *BRCA1* potentiated the DNA damage caused by ERα signaling.

## DISCUSSION

Estrogens play a leading role in breast cancer risk and progression. Although women are exposed to estrogens from menarche to menopause, only 12% develop breast cancer suggesting the pathogenic actions of estrogen signaling varies among individuals. Genetic modifiers of ERα signaling may be among the more than 300 polymorphic loci linked to breast cancer risk [15,17,21,75]. The inability of present culture methods to preserve expression of the endogenous ERα in normal breast epithelial cells has been a substantial hurdle in Previous efforts to recover ERα signaling in HBECs using constitutive expression showed limited success [76–85]. As an alternate approach, we used conditional expression of ERα. The lentiviral vector had minimal basal expression preventing detrimental effects of ERα signaling, but generated levels of ERα similar to MCF7 breast cancer cells using 100ng/mL of doxycycline **(Figure 1B)**. The use of pooled populations of cells representing diverse integration sites minimizes the potential for artifacts due to clonal selection. For studying transcriptional responses, the 100 ng/mL concentration of doxycycline was used to avoid possible metabolic effects [86] and 10nM E2 approximates physiologic levels found during mid-pregnancy [87,88]. These conditions allowed ligand-dependent transcriptional activation of endogenous target genes in all the cell lines **(Figure 2E)** as well as increased proliferation in 3 of the cell lines **(Figure 3)**. Conditioned media from 2 cell lines were used to determine if secreted growth factors were responsible for proliferative responses. The conditioned media failed to induce proliferation in the parental cell lines lacking ERα indicating that secreted factors were not sufficient to induce proliferation **(Figure 4)**. Therefore, intrinsic signaling mediated by ERα is necessary for ERα-mediated proliferation in these cells.

RNA-Seq was used to evaluate the transcriptome induced by ERα signaling in the 4 HBEC-*ESR1* cell lines. A total of 682 E2-responsive genes were identified among the 4 HBEC-*ESR1* lines conditionally expressing ERα **(Figure 5C)**. These represent a core set of target genes and are also regulated similarly by E2 in the ERα+ MCF7 and T47D breast cancer cell lines **(Figure 5D&E)**. Therefore, conditional expression of ERα was able to recover a large portion of the ERα transcriptome.

While core responses are observed in the cell lines, each showed distinctive overall patterns of ERα-mediated transcriptional responses. The differences in estrogen responsive transcriptomes are also evident in the MCF7 and T47D breast cancer cell lines which shared only 17.8% of the genes in common **(Figure 5D)**. The EstroGene database [89] compiles transcriptional responses in ERα+ breast cancer cell lines and shows similarly diverse responses to E2 among ERα+ breast cancer cell lines. This is, in part, attributable to the genomic alterations and differences in ploidy among the cancer cells. In contrast, the HBEC-*ESR1* cells have largely was overlap between 43% of estrogen-responsive genes in HBEC-*ESR1* cell lines and the breast cancer cell lines (MCF7, T47D) confirming the recovery of a large portion of the estrogen-responsive transcriptome. Hierarchical clustering of the transcriptional responses in HBEC-*ESR1* cells show consistent induction of genes with Dox+E2 which are also downregulated with Dox+ICI **(Figure 5E)**. In addition to these shared responses to ERα-mediated signaling, the HBEC-*ESR1* cell lines each had unique transcriptional profiles suggesting “private patterns”. Unique patterns of transcriptional responses to E2 was also observed in isolates of normal breast epithelial cells and differed substantially when compared to those or ERα+ breast cancers [33]. In this study, the luminal epithelium enriched by FACS were analyzed for acute transcriptional responses in 5 donors and from ERα+ breast cancers. This emphasizes a need for tools to understand how transcriptional responses mediated by ERα in normal breast tissues as well as the genetic and epigenetic differences among individuals that can alter the consequences of exposures to endogenous estrogens and environmental xenoestrogens. The inducible ERα system can allow identification of “private patterns” of ERα binding and transcriptional responses among individuals which may mediate differences in risk of breast cancer.

How E2 promotes breast cancer remains a significant debate. Direct mutagenesis can occur but requires supraphysiologic concentrations of E2 and the carcinogenic effect would not be restricted to breast tissue [5,6]. Recent work has highlighted the role of ERα-mediated DSBs. Co-transcriptional R-loops provide a mechanism for the DNA damage [9]. Xenoestrogens were also shown to increase DSBs in breast cancer cell lines and in mammary glands of mice [3]. Consistent with these observations, ERα-expressing HBECs stimulated with E2 resulted in enrichment of genes involved in DNA repair. Levels of nuclear γH2AX foci provide a surrogate to quantify DSBs in the cell lines. Both baseline and E2-induced DSBs varied among the cell lines **(Figure 6B)**. The MCF10A-*ESR1*, 76NTERT-*ESR1* and ME16C2-*ESR1* cell lines all showed significant increases in γH2AX foci when ERα signaling is induced by E2. This is, in part, due to cells entering S-phase but was also evident in the G1 population of cells **(Figure 6D)** suggesting that ERα signaling can be mutagenic. However, all women are exposed to E2 from menarche through menopause, but breast cancer is diagnosed in fewer than 12% of women This is most evident in women carrying germline highly penetrant pathogenic variants in genes involved in homology-directed DSB repair. Therefore, we used small molecule inhibitors of HR and NHEJ pathways to impair DSB repair mechanisms (**Figure 7)**. Inhibition of HR with Mirin amplified the DSBs detected using γH2AX compared to the expression of ERα alone (Dox+Control treated) in both cell lines tested. Inhibition of NHEJ also significantly increased γH2AX in ME16C2-*ESR1* cells and to a lesser extent in MCF10A-*ESR1* cells. The increase in DSBs by E2 was also observed in mammary glands of BALB/c mice and was further increased by disruption of repair by NHEJ in mice lacking *Prkdc* and resulted in rapid development of premalignant lesions [4]. The BALB/c and C57BL/6 mice differ in susceptibility to mammary tumors which was genetically linked to a reliance on error-prone repair of DSBs via alternative end-joining in BALB/c [90]. Therefore, polymorphisms disrupting DSB repair pathways can amplify differences in fidelity of DNA replication and repair mechanisms causing variation consequences of estrogen exposure on breast cancer susceptibility among individuals.

Germline mutations in *BRCA1, BRCA2* and *PALB2* affect DSB repair through homologous recombination and preferentially predispose to breast cancer. Therefore, effects of ERα signaling were compared in isogenic MCF10A cells with either wildtype or a heterozygous frameshift mutation in *BRCA1* **(Figure 7D&E)**. Induction of ERα signaling resulted in significantly more DSBs in MCF10A-*BRCA1^185del/+^-ESR1* compared to MCF10A-*ESR1* indicating that haploinsufficiency in DSB repair can exacerbate the DNA damage in normal luminal breast epithelial cells which express ERα. While ERα+ is a biomarker of good prognosis in the general population, however, the overall survival is significantly worse for those with ERα+ breast cancers in carriers of germline mutations in *BRCA1/2* compared to carriers with TNBCs [91–95]. These observations suggest a mechanism where defects in DSB repair are amplified in tissues where ERα signaling is most active resulting in a preferential susceptibility to breast cancer.

These results demonstrate the utility of inducible expression of ERα to reconstruct transcriptional responses in immortalized breast epithelial cells. The immortalized HBEC-*ESR1* cells express ERα that is inducible with doxycycline and demonstrate ligand-dependent E2 responses such as gene expression, proliferation and ERα-mediated DNA damage. The methods identify a core set of genes that are responsive in both HBEC-*ESR1* cells and breast cancer cell lines. However, the results also reveal variation in estrogen-responsive transcriptomes among individuals. As the cells are largely euploid, the variation in ERα-mediated transcription appears to reflect differences in genetic polymorphisms as well as epigenetic signatures among individual donors. The method of ERα expression can be especially useful in extending analyses to HBECs from genetically diverse donors based on polygenic risk scores and inherited risk alleles to identify genetic polymorphisms that alter ERα signaling and are linked to breast cancer risk. While the availability of immortalized normal breast epithelial cells remains a limitation, new efforts to establish these cells reliably are showing promise [96].

## ABBREVIATION

E2: Estrogen, 17-β-estradiol
ER: Estrogen Receptor
HBEC: Human breast epithelial cells
DSB: double strand break
DSBR: double strand break repair
pIND-*ESR1*: Inducible *ESR1* construct in pINDUCER vector backbone
ICI: ICI 182,780, also known as Fulvestrant

