## Supplementary data for "Inducible estrogen receptor alpha in normal breast epithelial cells demonstrate estrogen receptor-dependent DNA damage"

### Supplemental Figures

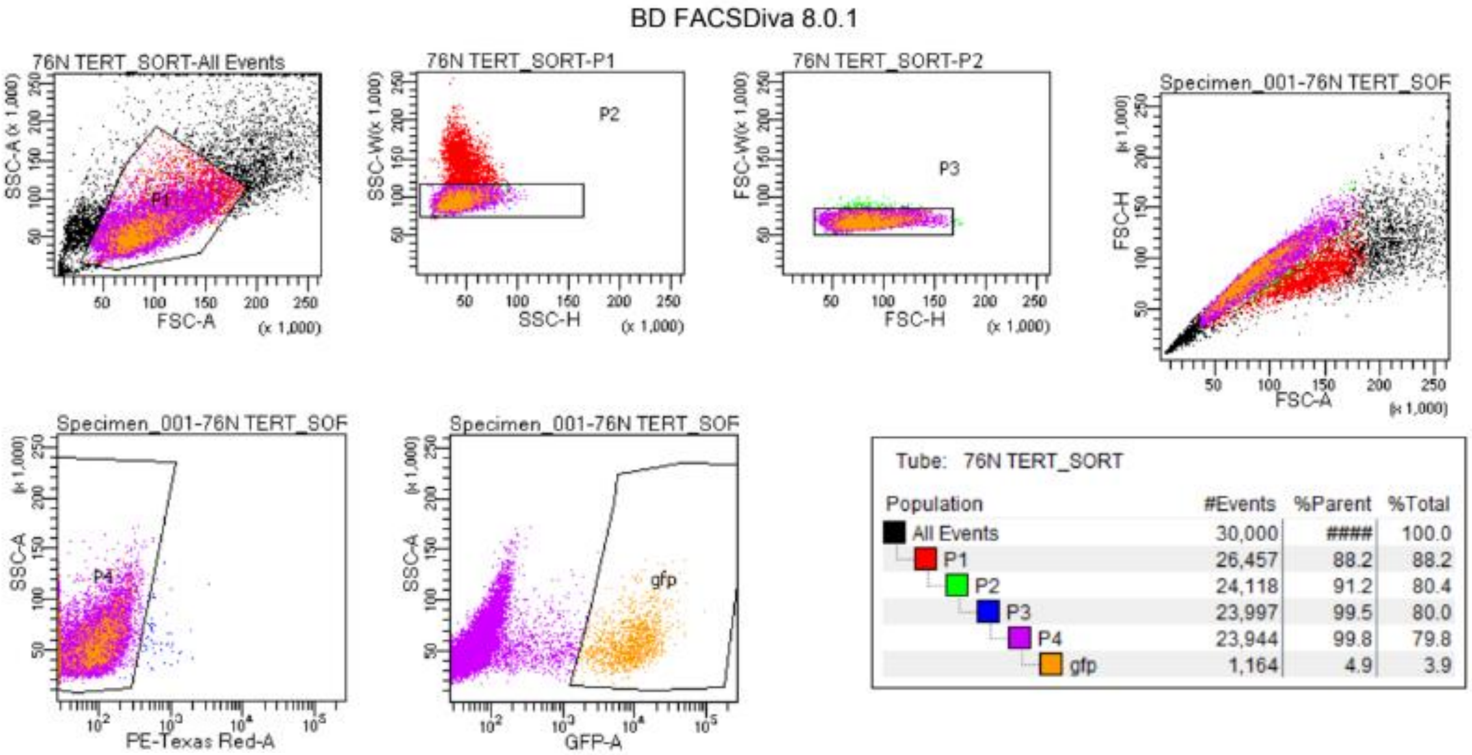


**Supplemental Figure 1: FACS selection of 76N TERT-*ESR1* cells**

76N TERT cells infected with pIND-*ESR1* were selected for GFP expression using FACS. Side scatter area (SSC-A) and forward scatter area (FSC-A) were used to isolate viable cells, SSC width (W) and height (H) were used to remove doublets, and FSC-W and FSC-H confirmed selection of single cells. PE-Texas Red showed minimal autofluorescence. Parental 76N TERT cells were used as a negative control to set background autofluorescence. Approximately 5% of cells were GFP positive and are considered the new 76N TERT-*ESR1* cell line.

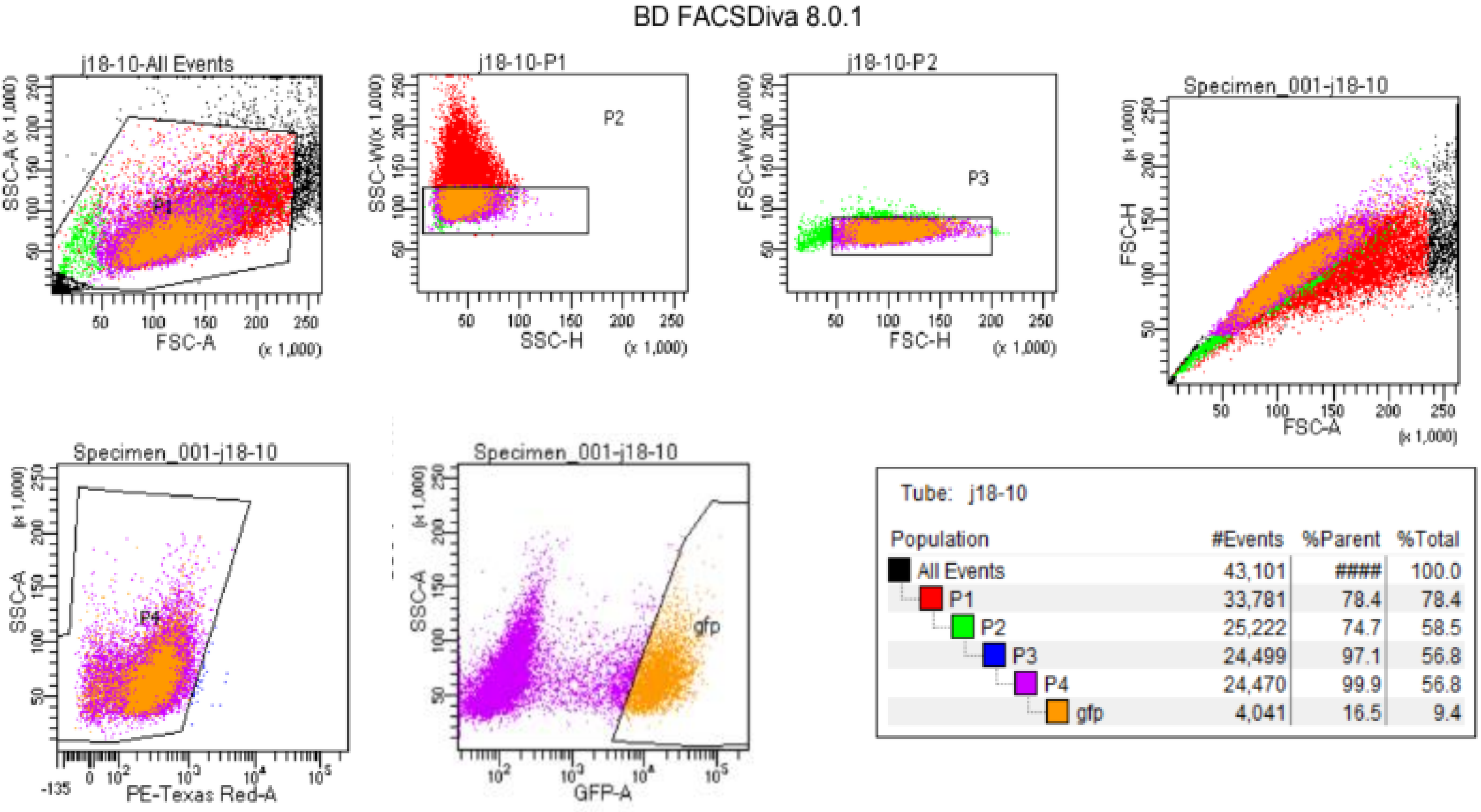


**Supplemental Figure 2: FACS selection of MCF10A-*ESR1* cells**

MCF10A cells infected with pIND-*ESR1* were selected for GFP expression using FACS. Side scatter area (SSC-A) and forward scatter area (FSC-A) were used to isolate viable cells, SSC width (W) and height (H) were used to remove doublets, and FSC-W and FSC-H confirmed selection of single cells. PE-Texas Red showed minimal autofluorescence. Parental MCF10A cells were used as a negative control to set background autofluorescence. Approximately 17% of cells were GFP positive and are considered the new MCF10A-*ESR1* cell line.

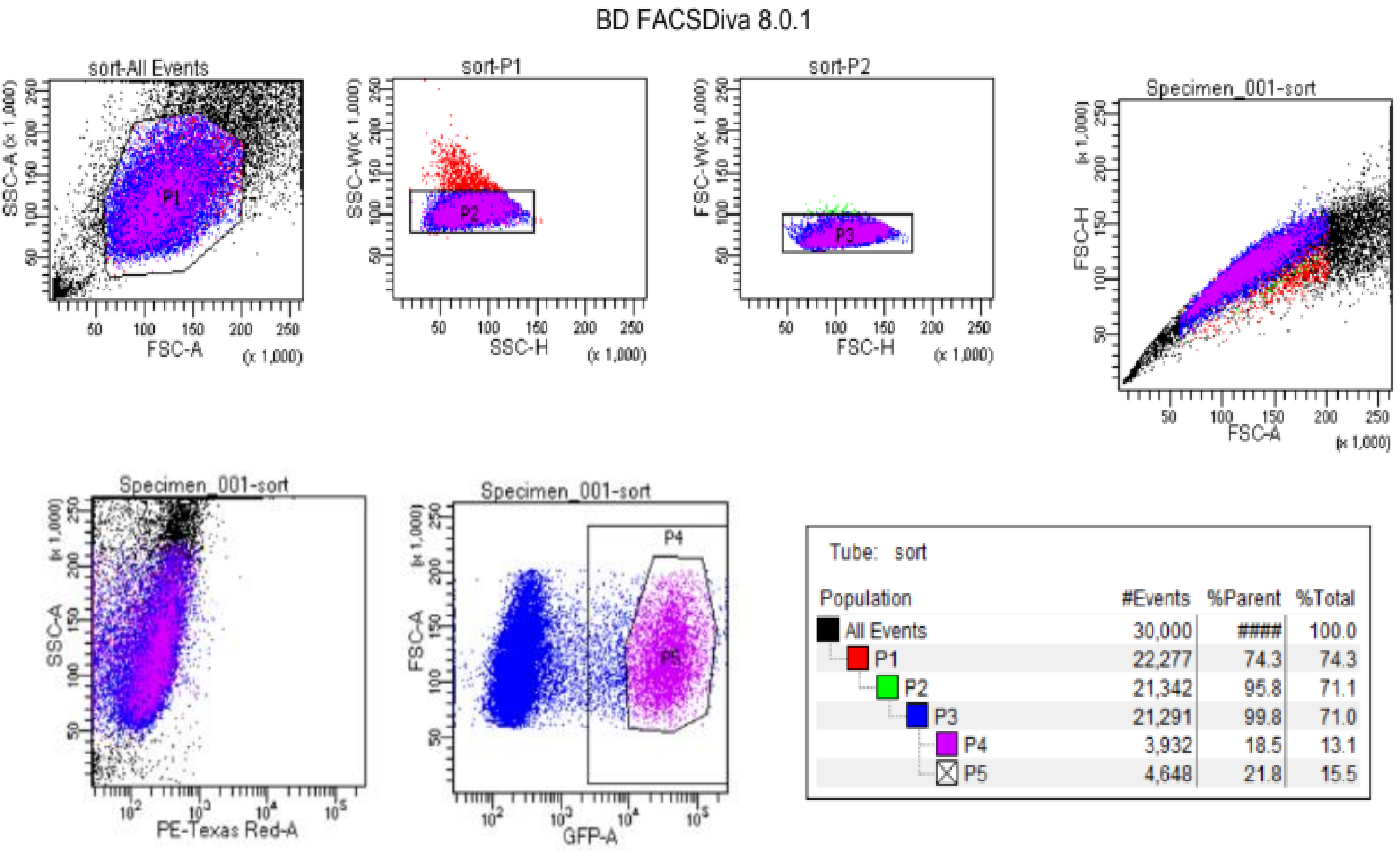


| Population | #Events | %Parent | %Total |
|---|---|---|---|
| All Events | 30,000 | #### | 100.0 |
| P1 | 22,277 | 74.3 | 74.3 |
| P2 | 21,342 | 95.8 | 71.1 |
| P3 | 21,291 | 99.8 | 71.0 |
| P4 | 3,932 | 18.5 | 13.1 |
| P5 | 4,648 | 21.8 | 15.5 |

**Supplemental Figure 3: FACS selection of HME-CC-*ESR1* cells**

HME-CC cells infected with pIND-*ESR1* were selected for GFP expression using FACS. Side scatter area (SSC-A) and forward scatter area (FSC-A) were used to isolate viable cells, SSC width (W) and height (H) were used to remove doublets, and FSC-W and FSC-H confirmed selection of single cells. PE-Texas Red showed minimal autofluorescence. Parental HME-CC cells were used as a negative control to set background autofluorescence. Approximately 22% of cells were GFP positive and are considered the new HME-CC-*ESR1* cell line.

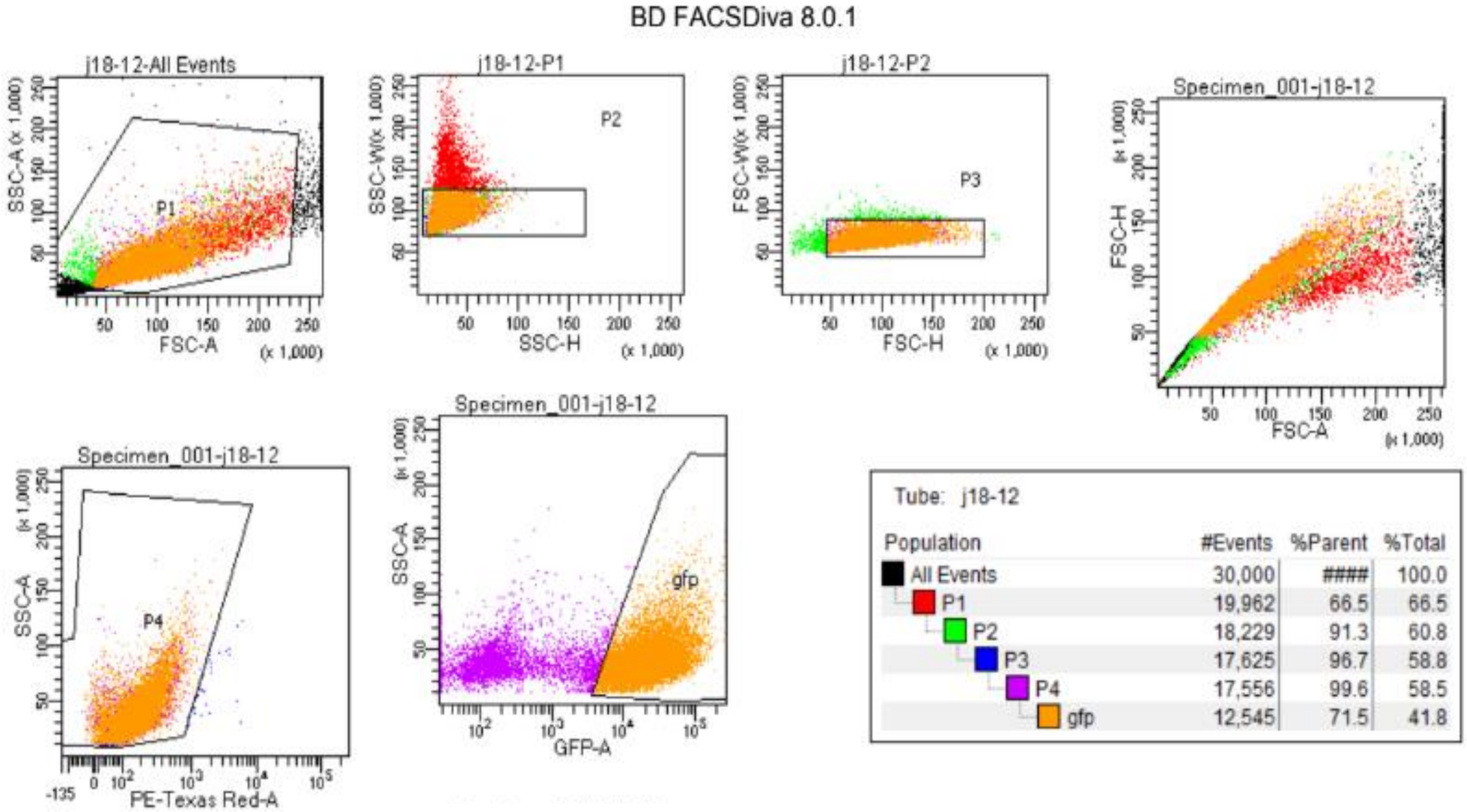


| Population | #Events | %Parent | %Total |
|---|---|---|---|
| All Events | 30,000 | #### | 100.0 |
| P1 | 19,962 | 66.5 | 66.5 |
| P2 | 18,229 | 91.3 | 60.8 |
| P3 | 17,625 | 96.7 | 58.8 |
| P4 | 17,556 | 99.6 | 58.5 |
| gfp | 12,545 | 71.5 | 41.8 |

**Supplemental Figure 4: FACS selection of ME16C2-*ESR1* cells**

ME16C2 cells infected with pIND-*ESR1* were selected for GFP expression using FACS. Side scatter area (SSC-A) and forward scatter area (FSC-A) were used to isolate viable cells, SSC width (W) and height (H) were used to remove doublets, and FSC-W and FSC-H confirmed selection of single cells. PE-Texas Red showed minimal autofluorescence. Parental ME16C2 cells were used as a negative control to set background autofluorescence. Approximately 72% of cells were GFP positive and are considered the new ME16C2-*ESR1* cell line.

A

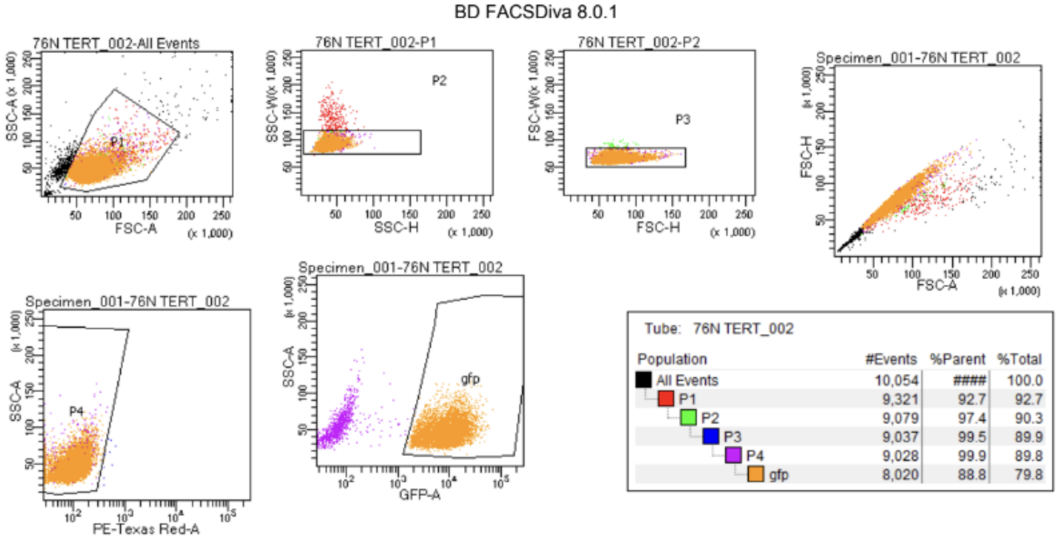


Tube: 76N TERT_002

| Population | #Events | %Parent | %Total |
|---|---|---|---|
| All Events | 10,054 | #### | 100.0 |
| P1 | 9,321 | 92.7 | 92.7 |
| P2 | 9,079 | 97.4 | 90.3 |
| P3 | 9,037 | 99.5 | 89.9 |
| P4 | 9,028 | 99.9 | 89.8 |
| gfp | 8,020 | 88.8 | 79.8 |

B

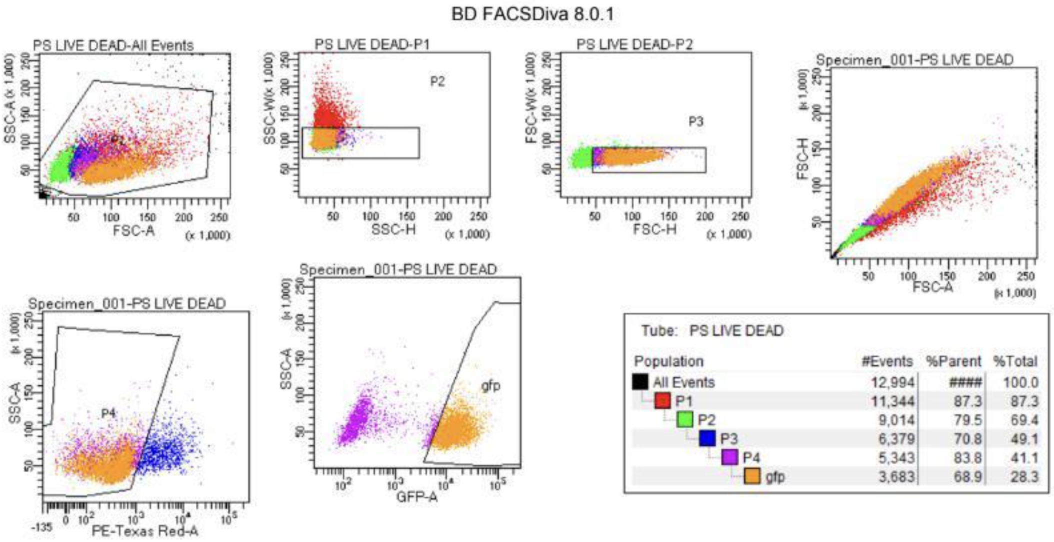


Tube: PS LIVE DEAD

| Population | #Events | %Parent | %Total |
|---|---|---|---|
| All Events | 12,994 | #### | 100.0 |
| P1 | 11,344 | 87.3 | 87.3 |
| P2 | 9,014 | 79.5 | 69.4 |
| P3 | 6,379 | 70.8 | 49.1 |
| P4 | 5,343 | 83.8 | 41.1 |
| gfp | 3,683 | 68.9 | 28.3 |

C

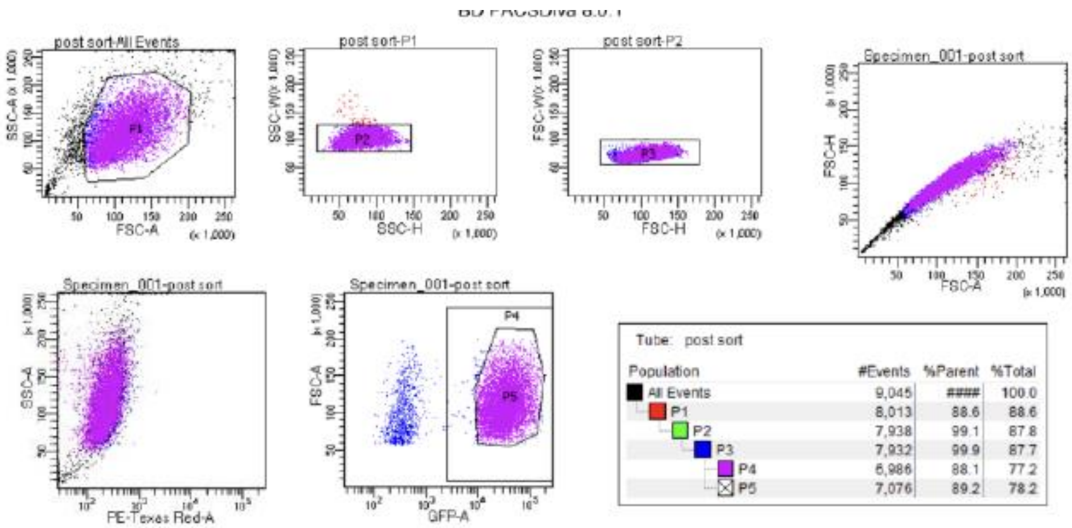


Tube: post sort

| Population | #Events | %Parent | %Total |
|---|---|---|---|
| All Events | 9,045 | #### | 100.0 |
| P1 | 8,013 | 88.6 | 88.6 |
| P2 | 7,938 | 99.1 | 87.8 |
| P3 | 7,932 | 99.9 | 87.7 |
| P4 | 6,986 | 88.1 | 77.2 |
| P5 | 7,076 | 89.2 | 78.2 |

**D**

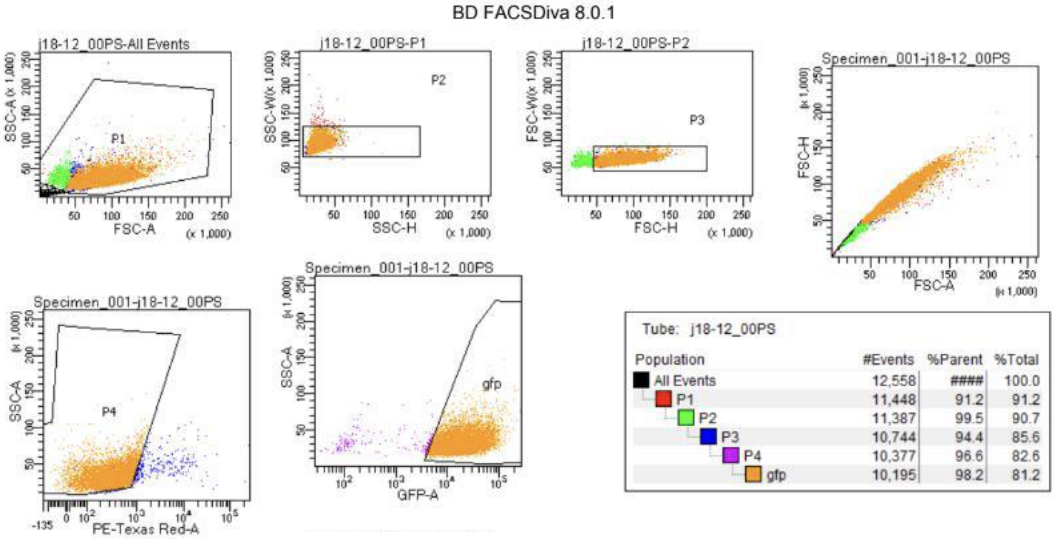


**Supplemental Figure 5: Proportion of GFP+ HBEC-*ESR1* cells post-FACS sorting.**
After FACS for GFP expressing each HBEC cells, an aliquot was checked for purity. Side scatter area (SSC-A) and forward scatter area (FSC-A) were used to isolate viable cells, SSC width (W) and height (H) were used to remove doublets, FSC-W and FSC-H confirmed selection of single cells, and PE-Texas Red was used to detect autofluorescence. **A**) Approximately 90% of 76-N-TERT-*ESR1* cell lines are GFP+, **B**) 70% of the sorted MCF10A-*ESR1* population expresses GFP, **C**) 90% of HMEC-CC-*ESR1* cells express GFP and **D**) approximately 72% of ME16C2 cells were GFP positive

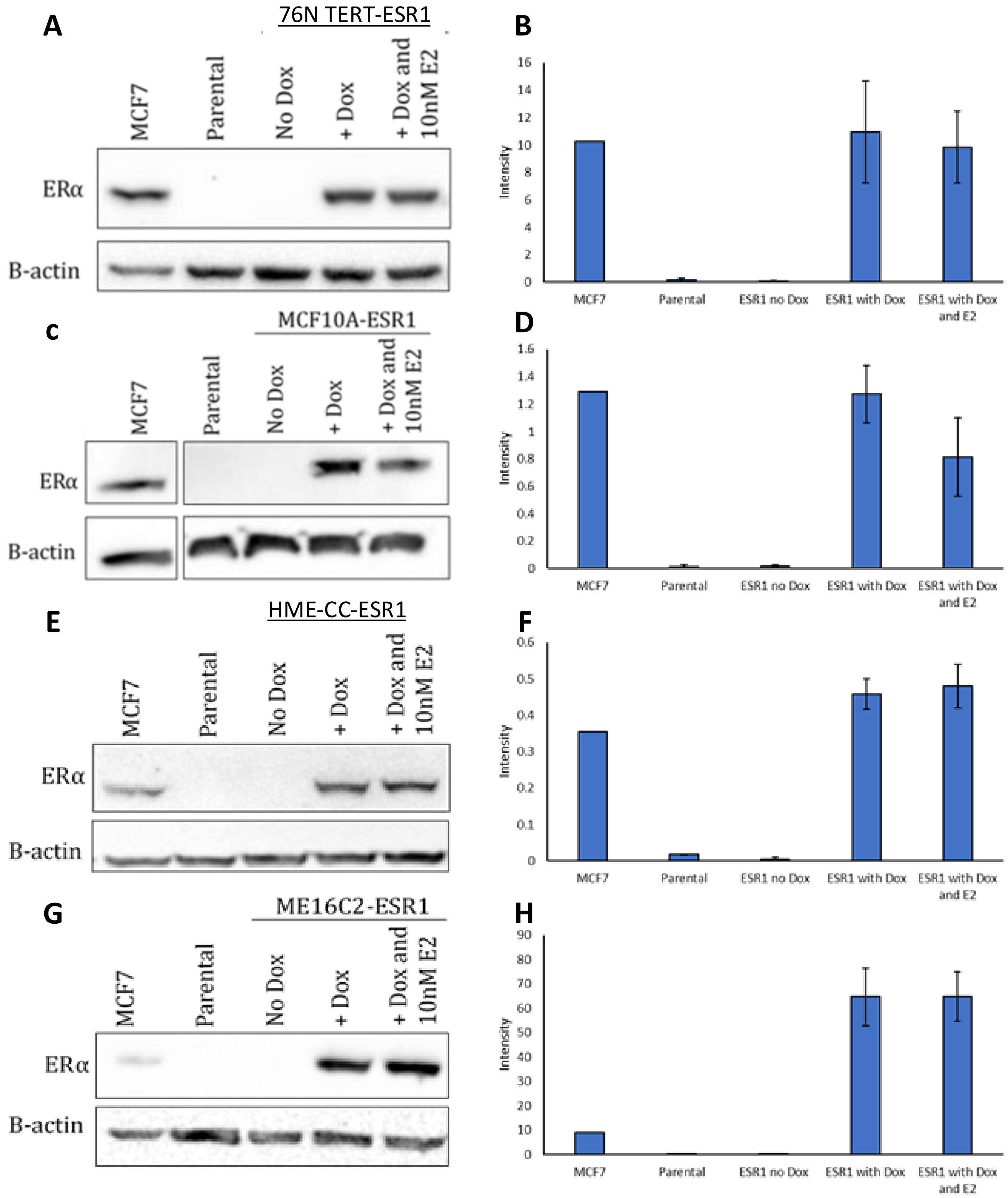


**Supplemental Figure 6: ERα protein expression in HBEC-*ESR1***

Detection and quantification ERα protein levels in 76N TERT-*ESR1* (**A, B**), MCF10A-*ESR1* (**C, D**), HME-CC-*ESR1* (**E, F**), and ME16C2-*ESR1* (**G, H**) cell lines. MCF7 cells were included as a positive control for ERα, and each parental HBEC line included as a negative control. β-actin was detected as a protein loading control. HBEC-*ESR1* lines were treated with 0ng/mL (No Dox) or 100ng/mL doxycycline (+Dox) for 48 hours and either control or 10nM E2 media for 24 hours (+Dox and 10nmE2) before lysate collection. Quantification of ERα protein was performed for each cell line by normalizing to β-actin expression. Error bars indicate SEM.

| Target | Primer | Sequence |
|---|---|---|
| TFF1 | F | GATCCCTGCAGAAGTGTCTAAAA |
| | R | CCCCTGGTGCTTCTATCCTAA |
| GREB1 | F | ATC TTT TCC CAG CTG TAC CTG |
| | R | CTC CTT CTC CCT CTT CTT CCT |

**Supplemental Table 1: Primer Sequences used in qRT-PCR experiments.**
